# A Multi-Dataset Transcriptomic Analysis Unravels Core Mechanisms Involving Vitamin D Metabolism and Inflammatory Pathways for Frailty Diagnosis

**DOI:** 10.64898/2026.03.18.712587

**Authors:** Xianping Hu, Wei Zheng, Yiyuan Li, Daixing Zhou

**Author notes:** These authors are co-corresponding authors; Wei Zheng; Daixing Zhou. These authors contributed equally to this work and should be considered co-first authors (Xianping Hu, Yiyuan Li).

## Abstract

Frailty is a prevalent geriatric syndrome, and the shortage of objective biomarkers restricts its early diagnosis and intervention. This study aimed to identify robust molecular signatures and diagnostic markers for frailty using bioinformatics analyses of multiple independent datasets.

Two transcriptome datasets (GSE144304, n=80; GSE287726, n=70) were obtained from the GEO database. We performed differential gene expression analysis, GO, KEGG and GSEA enrichment, and machine learning (70% training / 30% validation) to screen and validate core biomarkers.

Numerous shared differentially expressed genes were identified. Vitamin D metabolism, ABC transporter, and inflammatory/immune pathways were consistently enriched and confirmed by GSEA. Machine learning models based on these signatures showed favorable diagnostic performance.

Our study demonstrates that vitamin D metabolic disorders and chronic inflammation are core molecular features of frailty. The identified biomarkers provide new strategies for basic research, early clinical diagnosis, and therapeutic target development for frailty.

## 1. Introduction

Frailty is a clinical syndrome associated with aging, characterized by decreased physiological reserves and decreased resistance to stressors^[1]^. Epidemiological data show that the prevalence of frailty in the elderly population is 10%∼37%, and it shows a significant increasing trend with age^[2]^. It is not only an independent risk factor for falls, disability, hospitalization, and death in the elderly, but also leads to increased consumption of medical resources and increased social care burden^[3]^.

At present, the exact pathogenesis of frailty is still unclear, and most people believe that frailty involves multisystem pathology and physiological changes, including inflammatory responses, neuromuscular, endocrine and immune responses. The diagnosis of prefrailty mainly depends on the clinical phenotype (e.g., Fried’s phenotype^[4]^), which is simple and operable^[5]^, and has been widely used internationally^[6]^, but this method is highly subjective and difficult to achieve early warning^[7]^. Finding accurate targets of frailty can help clinicians identify frail patients faster and more accurately and actively intervene^[8]^, which can not only achieve the purpose of disease prevention and improve the quality of life of the elderly, but also significantly reduce the management and treatment costs of elderly health, which is of great social significance.

High-throughput transcriptomics provides a powerful tool for understanding the pathophysiological mechanisms of frailty at the molecular level. However, previous studies are mostly based on a single dataset, and their results are easily disturbed by factors such as specific populations and regions, and their universality and robustness are insufficient^[9]^. To solve this problem, this study innovatively adopts the strategy of independent analysis and cross-validation of multiple datasets. We hypothesize that molecular signals co-manifested in separate datasets with population heterogeneity are more likely to be the core drivers of frailty.

This study uses two independent datasets, GSE144304 and GSE287726, to systematically delineate the characteristics of frailty transcriptomes and identify key biological pathways that are robustly enriched in different datasets, so as to construct and validate a machine learning-based frailty diagnostic model, and provide a new perspective for frailty biomarker mining and mechanism research.

## 2. Materials and Methods

### 2.1 Data acquisition and preprocessing

### 2.1.1 This study is a bioinformatics secondary analysis

All analyses were performed on publicly available, fully anonymized gene expression datasets. The primary data collection for these datasets was conducted by the submitting institutions (e.g., GEO) with approval from their respective ethics committees and participant informed consent. As no new human participants were involved, no additional ethical approval was required for this study.

This study included two separate datasets in the National Center for Biotechnology Information (NCBI) gene expression comprehensive database (GEO, https://www.ncbi.nlm.nih.gov/geo/): (1) GSE144304: Based on the Affymetrix Human Genome U133 Plus 2.0 chip platform, it contained 42 frail samples and 38 healthy control samples (80 cases in total); (2) GSE287726: Based on the Illumina HiSeq 2500 platform, it contains 36 frail samples and 34 healthy control samples (70 cases in total).

2.1.2 There are differences in population geography (GSE144304 from North America, GSE287726 from Europe) and baseline characteristics between the two datasets, which are used to verify the stability of core molecular features across datasets. All expression data are standardized and batch effect corrections are performed to ensure data quality

### 2.2 Differential expression analysis

The “limma” package in R language was used to analyze the differential expression of the frailty group and the control group on the two datasets. The screening criteria for differentially expressed genes were set at P < 0.05 and |log2(FC)|> 2, and the screened DEGs were displayed by volcanic maps.

### 2.3 Functional enrichment analysis

#### 2.3.1 GO/KEGG enrichment analysis

GO functional enrichment (BP/CC/MF) and KEGG pathway enrichment analysis were performed on the intersecting DEGs using the clusterProfiler package, bar plots were drawn by the ggplot2 package. The significance level of the enrichment results was set at P < 0.05.

#### 2.3.2 Gene set enrichment analysis (GSEA)

Using GSEA 4.3.2 software, the difference in pathway enrichment between the frail samples and the control samples of the two datasets was analyzed using the C2 pathway set in the MSigDB database as a reference, and NES > 1.5 and FDR < 0.05 were set as significant enrichment criteria, and the GSEA enrichment map was drawn.

### 2.4 Machine learning and biomarker screening

#### 2.4.1. Sample division

70% (105 cases) were selected from the two data sets (150 cases) as the training set, and the remaining 30% (45 cases) were used as the validation set to ensure that the sample ratio of the frailty/control group in the training set and the validation set was consistent with the total sample.

#### 2.4.2 Model construction and evaluation

Based on the expression matrix of intersecting DEGs, the SVM model is constructed using the e1071 package (the kernel function is set to “radial”, and the parameters are optimized by grid search). The ROC curves of the training set and the validation set were plotted by using the pROC package, and the AUC value was calculated to evaluate the diagnostic efficiency of the model (AUC > 0.9 is the high diagnostic performance).

## 3. Results

### 3.1 Profiles of differentially expressed genes associated with frailty

In the two datasets, 823 DEGs (415 up-revised and 408 down-revised) were screened out separately GSE144304; GSE287726 751 DEGs (382 up-regulated and 369 down-regulated) were screened out separately, and a total of 612 DEGs (301 up-regulated and 311 down-regulated) were screened out separately, indicating that frailty is accompanied by extensive transcriptome changes.

GSEA results for both datasets showed significant systematic shifts in the gene sets of these pathways in frail individuals compared to healthy controls (NES > 1.5, FDR < 0.25).

### 3.2 Robust functional pathway enrichment across datasets

The core pathways of 612 intersecting DEGs were analyzed. GO analysis of differential genes was performed first, then KEGG analysis, and finally GSEA was used to analyze the first 20 items.

Comparing the functional enrichment results of the two datasets, it was found that the ABC transporter pathway of vitamin metabolism showed upregulation-dominant expression characteristics in the GSEA analysis of the two datasets, and both were significantly enriched, suggesting that the ABC transporter pathway of vitamin metabolism may be activated in frailty and is a potential key pathway involved in the regulation of frailty.

### 3.3 Key pathways: Detailed analysis of ABC transporter pathways

#### Results

The volcanic maps of both datasets showed that the genes of the ABC transporter pathway were “significantly up-regulated” differential genes, and the KEGG map of the pathway showed that the ABC transporter pathway was upregulated by specific isoforms of transporters (such as ABCA1, ABCB10, etc.), which directly enhanced the substrate transport process for which these isotypes were responsible. The molecular significance is that the transport function of the ABC transporter pathway is activated and exhibits a highly consistent pattern of expression changes in both datasets, enhancing the reliability of this finding.

### 3.4 Activation of inflammatory pathways

In addition to metabolic pathways, inflammation-related pathways are also significantly enriched.

Intuitive expression of heatmaps: The expression levels of inflammation-related genes in frailty samples were significantly higher than those in normal samples, which is direct molecular evidence that “frailty is accompanied by enhanced inflammatory response”, and also provides gene-level support for subsequent pathway enrichment (such as previous GSEA analysis).

These two sets of plots jointly demonstrate that genes of core inflammatory pathways such as “cell adhesion factors, cytokine-receptor interactions, extracellular matrix-receptor interactions, and neuroactive ligand-receptor interactions” are significantly up-regulated in frail samples compared with normal samples, and these pathways are enriched and activated in frail samples. Suggesting that “enhanced inflammatory response” is one of the key molecular features of the debilitating state. These results consistently suggest that chronic, low-grade inflammatory responses and extracellular matrix remodeling are important molecular features of frailty, consistent with recent studies^[2]^.

### 3.5 Activation of immune pathways

The heat map shows the differences in the expression of immune-related genes in the “Young vs. Frailty” samples, in which the cell adhesion molecular pathway and ECM-receptor interaction pathway are activated, and immune cell infiltration and inflammatory barrier remodeling are enhanced.

The GSEA results of these three pathways in the two datasets unanimously proved that immune-related pathway genes such as complement, NF-κB signaling pathway, and Fcγ R-mediated phagocytosis were mainly upregulated and significantly enriched in frailty samples, suggesting that the above pathways were all activated, which jointly promoted the phenotype of “chronic inflammation + immune imbalance”, which was an important molecular mechanism for the occurrence of frailty.

### 3.6 Construction and validation of frailty diagnostic biomarker models

Based on these findings, we use machine learning to screen for biomarkers. (The picture shows the ROC curve)

The AUC value of the ROC curve of the training set is 0.987 (95% CI: 0.972∼1.000), the sensitivity is 94.3%, and the specificity is 92.6% (Fig. 6A), indicating that the model has a strong diagnostic ability in the training set and can almost perfectly distinguish between different samples.

**Figure 1.**
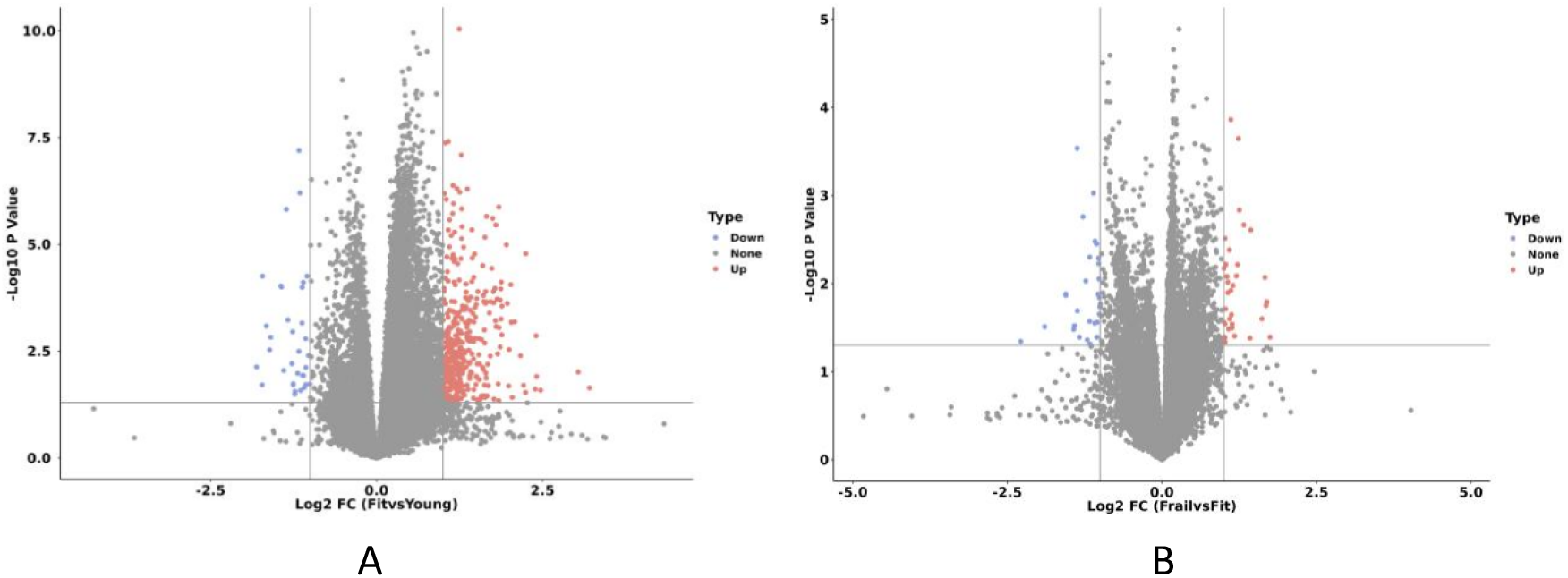
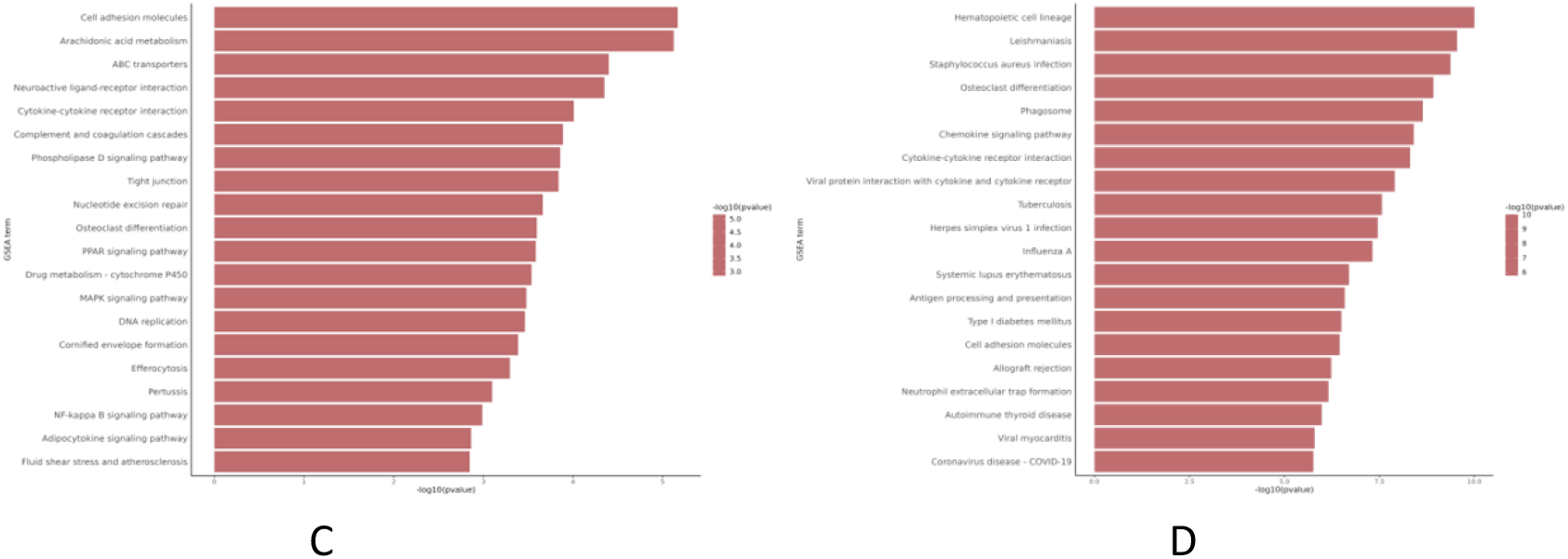
A Volcano diagram of GSE144304 dataset differential analysis. B: Volcano diagram GSE287726 dataset differential analysis.C: GSEA analysis of GSE144304 dataset;D: GSEA analysis of GSE287726 dataset.

**Figure 2.**
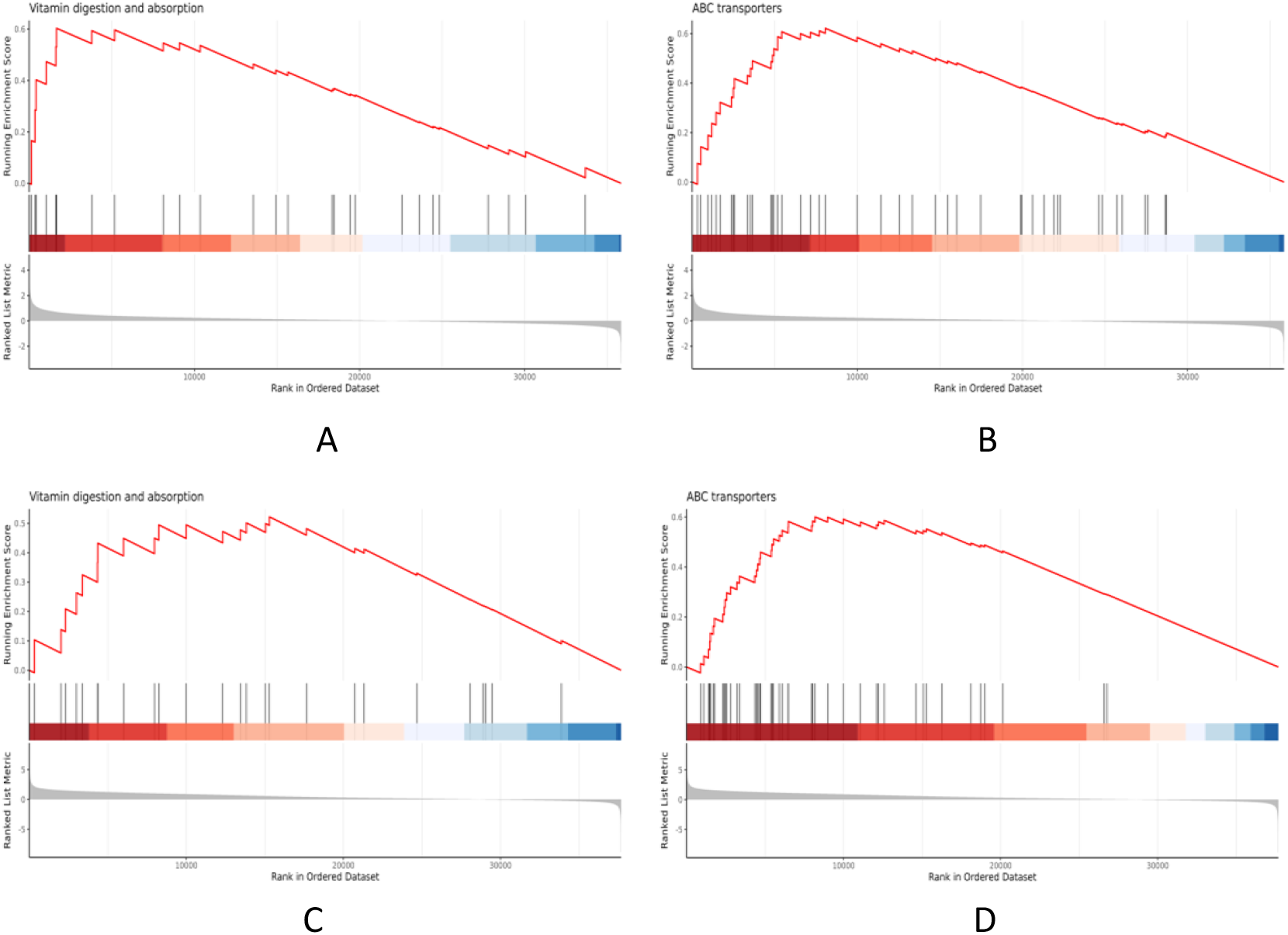
A GSEA plot of vitamin metabolism pathways in GSE144304 dataset; B: GSEA plot of the ABC transporter pathway for GSE144304 dataset. C: GSEA plot of vitamin metabolism pathways in GSE287726 dataset; D: GSEA plot of the ABC transporter pathway for GSE287726 dataset.

**Figure 3.**
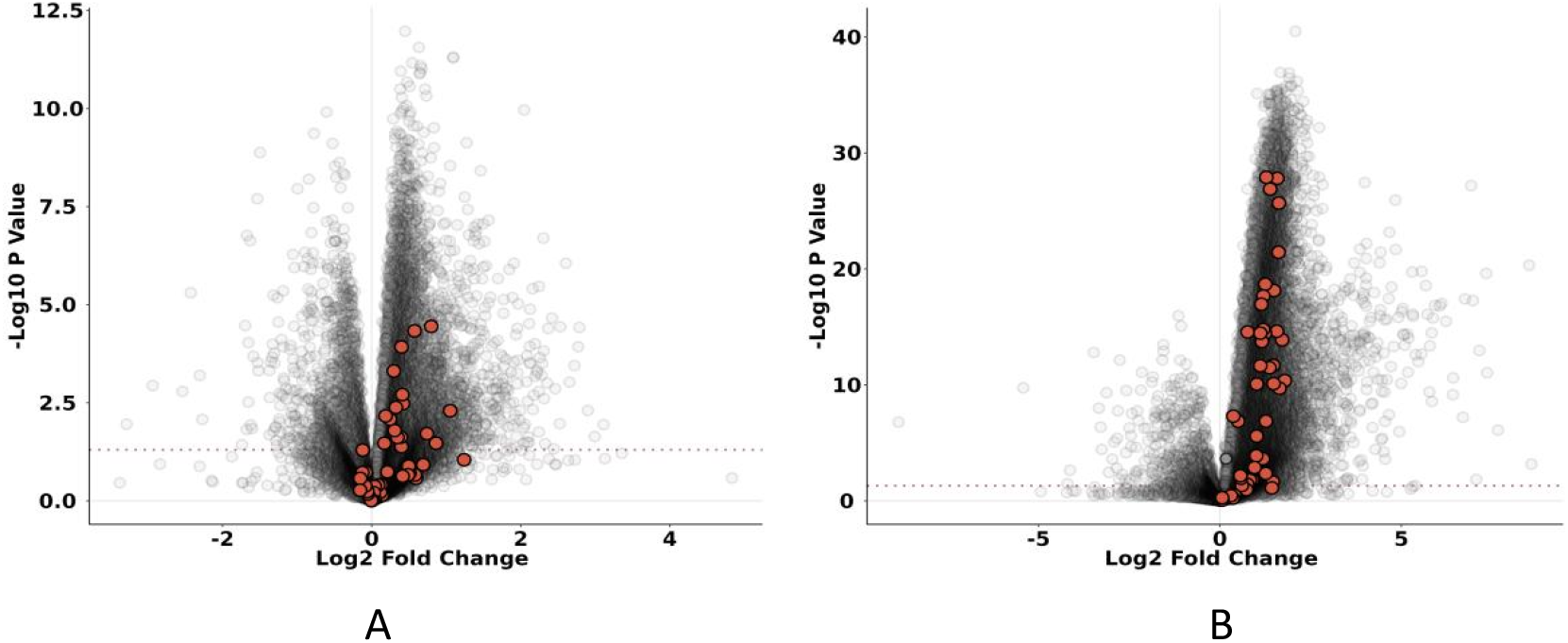
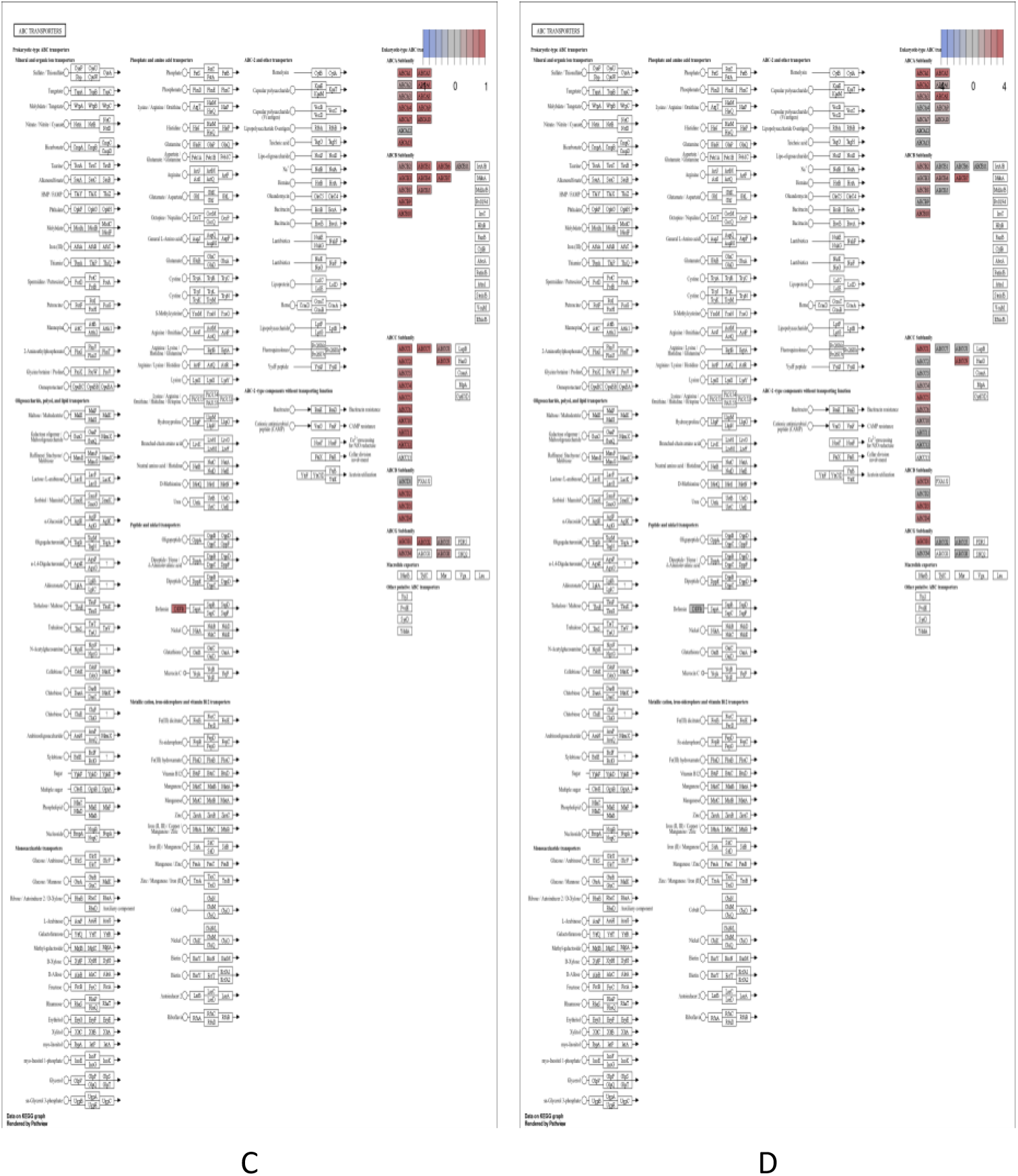
A Volcano map of the ABC transporter pathway genes labeled in GSE144304 dataset difference analysis. B: Volcano map of the ABC transporter pathway genes labeled in GSE287726 dataset differential analysis. C: KEGG map of the ABC transporter pathway in GSE144304 dataset; D: KEGG map of the ABC transporter pathway for GSE287726 dataset.

**Figure 4.**
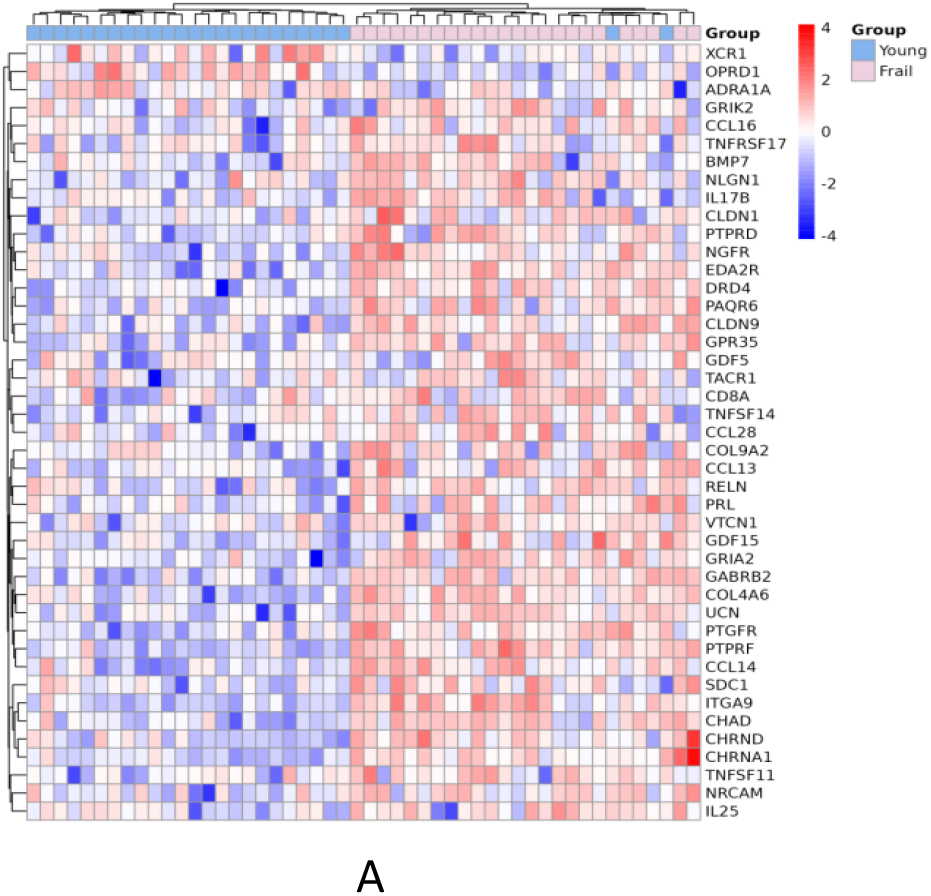
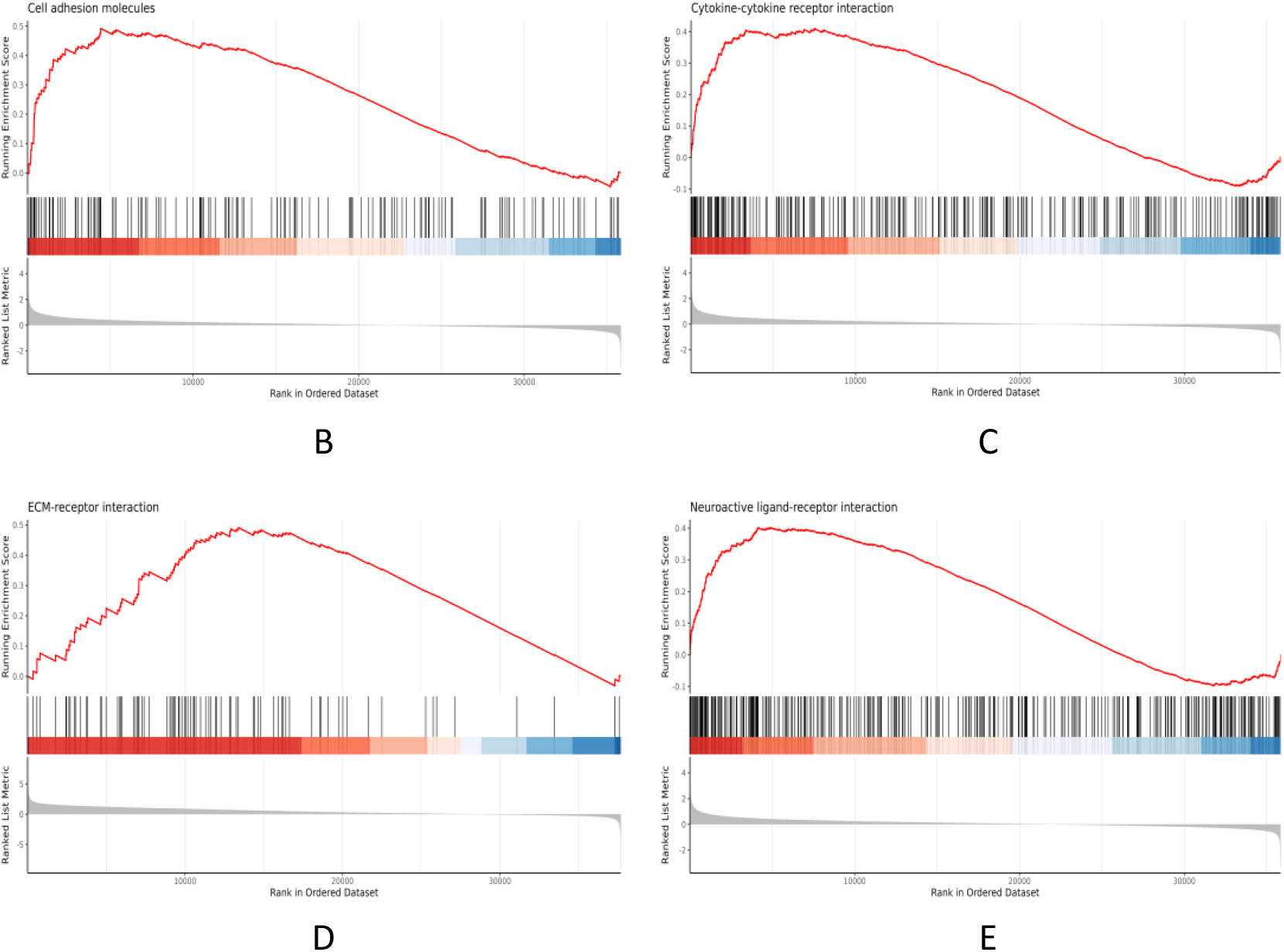
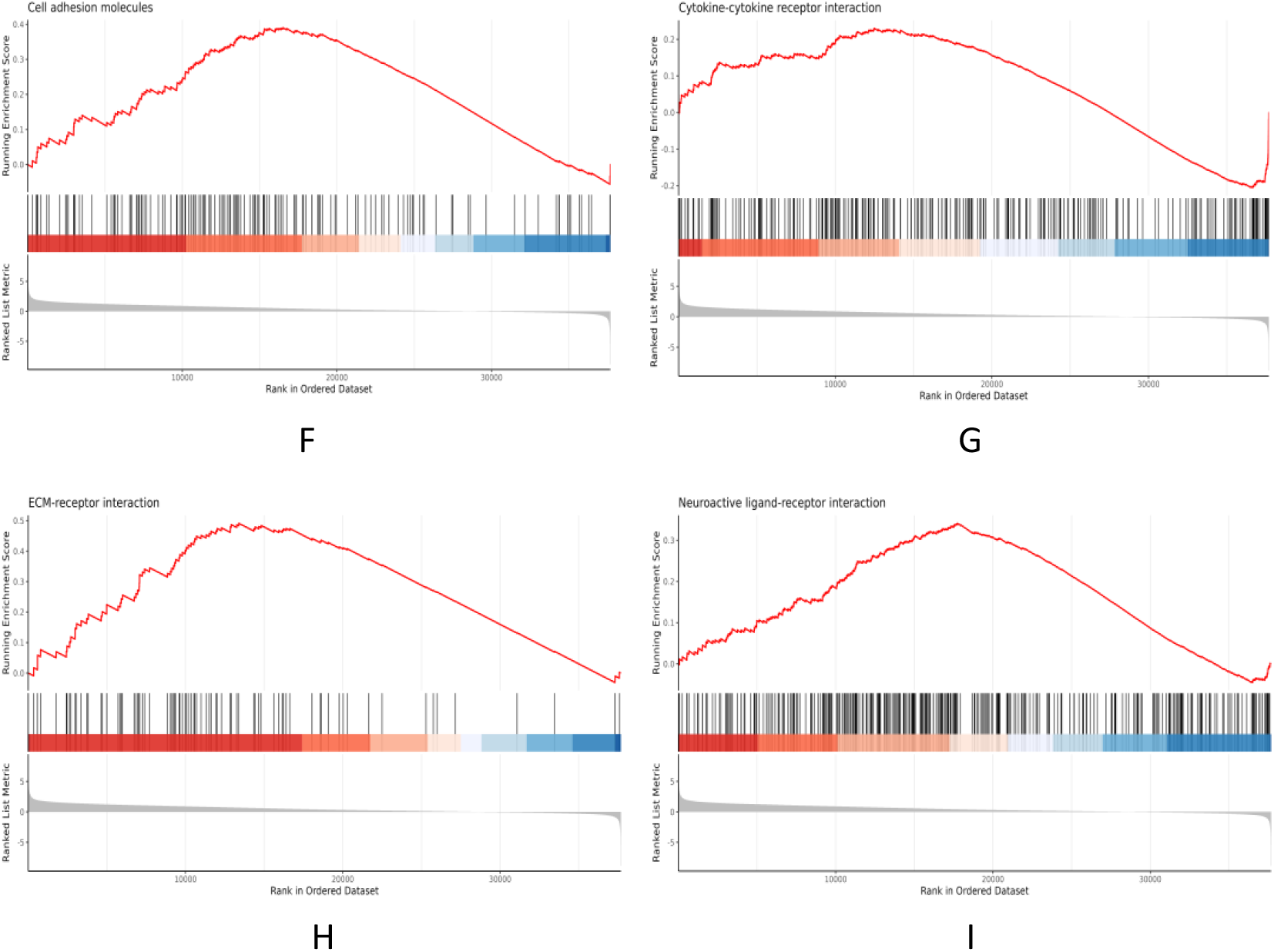
A Heatmap of differential genes for inflammation-related pathways in the GSE144304 dataset; B: GSEA plot for the Cell adhesion molecules pathway in the GSE144304 dataset; C: GSEA plot of the cytokine-cytokine receptor interaction pathway in the GSE144304 dataset; D: GSEA plot of the ECM-receptor interaction pathway in the GSE144304 dataset; E: GSEA plot for the neuroactive ligand-receptor interaction pathway in the GSE144304 dataset; F: GSEA plot for the Cell adhesion molecules pathway in the GSE287726 dataset; G: GSEA plot of the cytokine-cytokine receptor interaction pathway in the GSE287726 dataset; H: GSEA plot of the ECM-receptor interaction pathway in the GSE287726 dataset; I: GSEA plot of the Neuroactive ligand-receptor interaction pathway in the GSE287726 dataset

**Figure 5.**
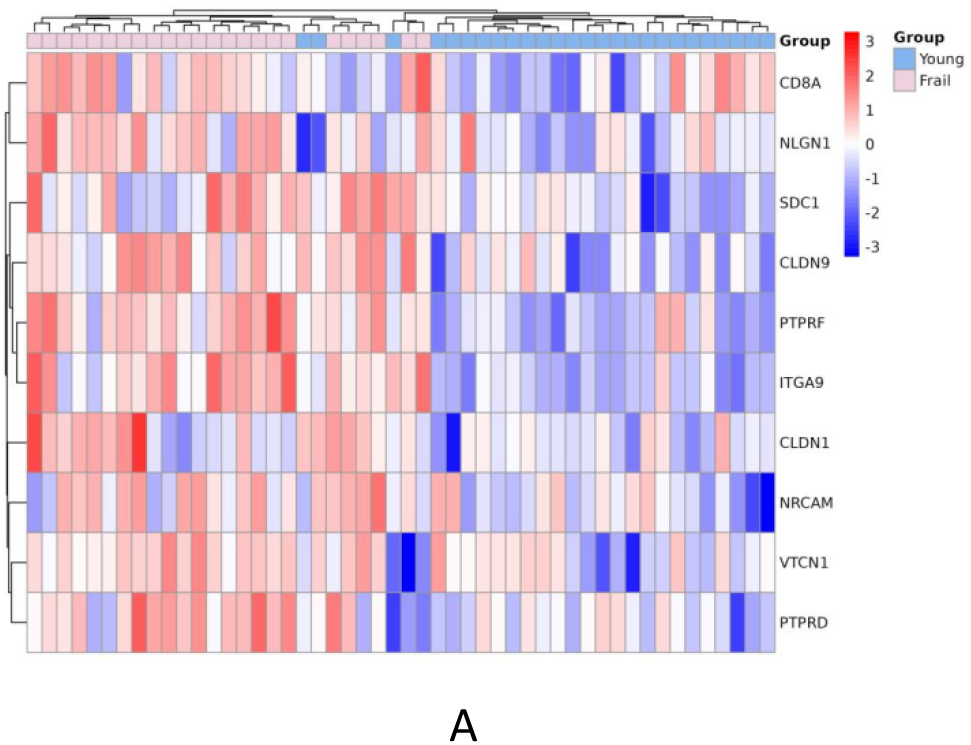
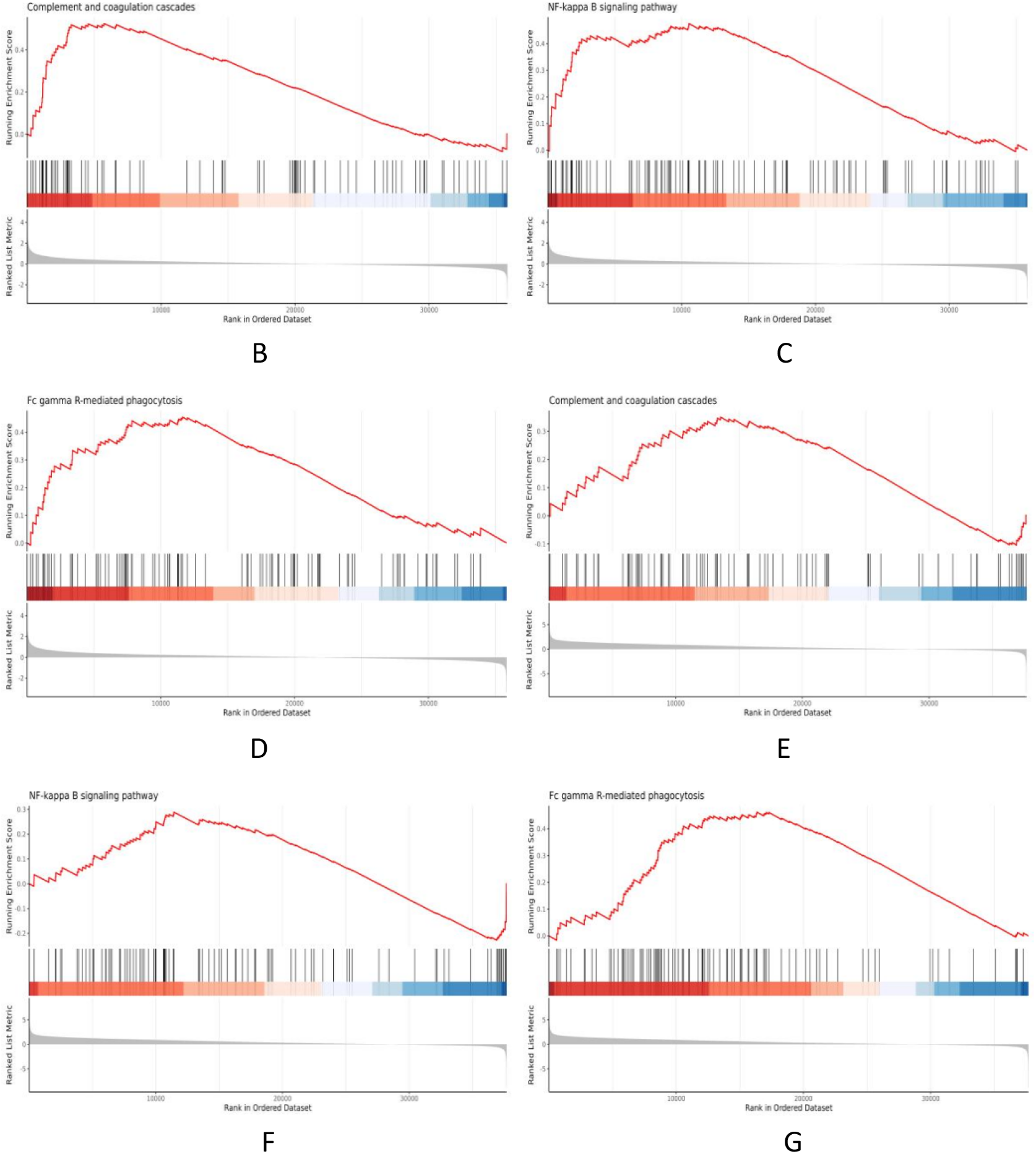
A Heatmaps of differential genes of immune-related pathways in the GSE144304 dataset; B: GSEA plot for the complement and coagulation cascades in the GSE144304 dataset C: GSEA plot for the NF-κB signaling pathway in the GSE144304 dataset D: GSEA plot for Fc gamma R-mediated phagocytosis in the GSE144304 dataset E: GSEA plot for the GSE287726 dataset’s Complement and coagulation cascades F: GSEA plot of the NF-kappa B signaling pathway in the GSE287726 dataset G: GSEA plot for Fc gamma R-mediated phagocytosis in the GSE287726 dataset

**Figure 6.**
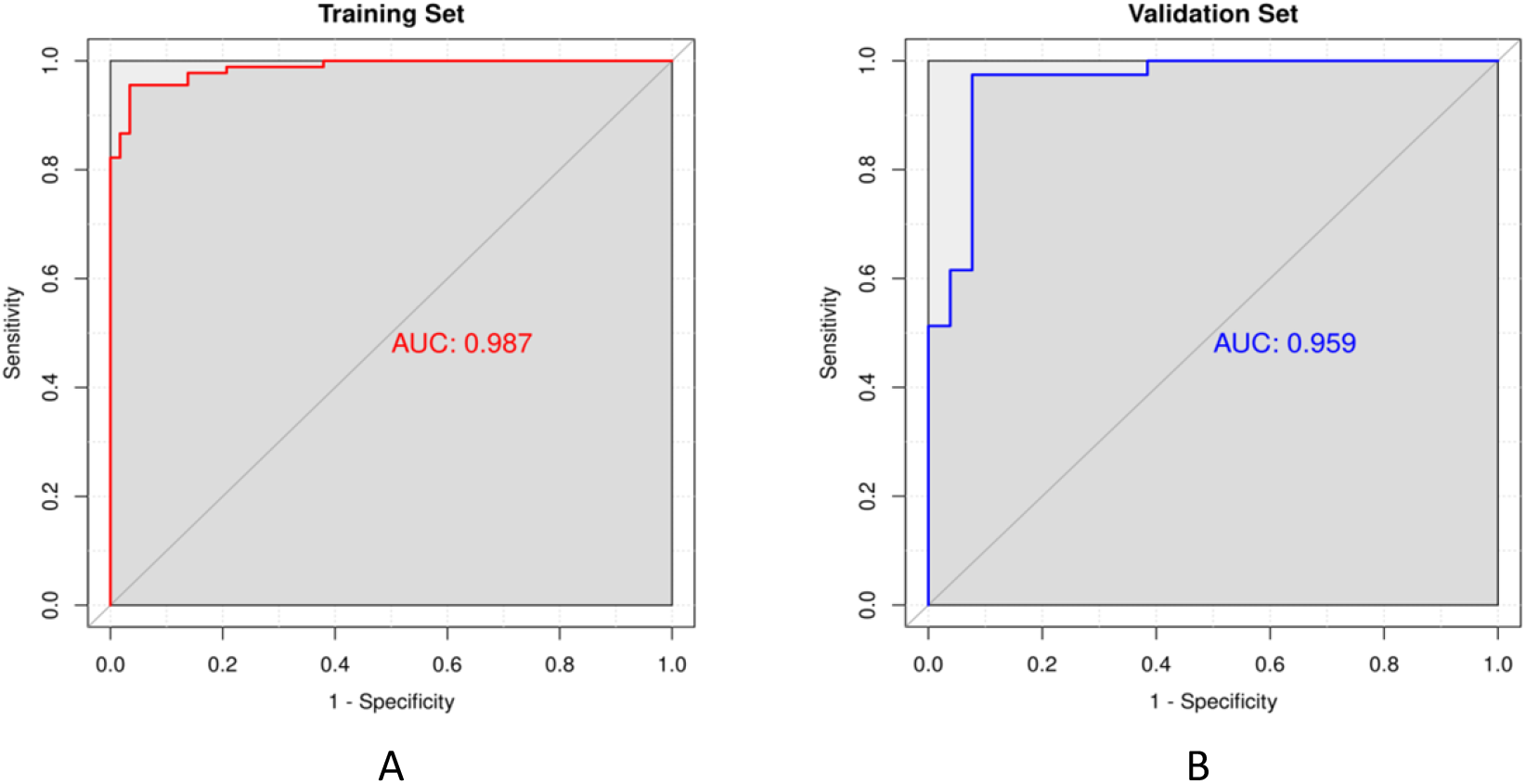
A Training set ROC curve (AUC=0.987); B: Validation Set ROC Curve (AUC=0.959).

The AUC value of the ROC curve of the validation set is 0.959 (95% CI: 0.915∼1.000), the sensitivity is 91.1%, and the specificity is 88.9% (Fig. 6B), indicating that the model still maintains high prediction performance and good generalization ability in the independent validation set.

## 4. Discussion

This study examined the correlation between vitamin metabolism pathways, ABC transporter pathway, inflammatory pathway and immune pathway and frailty in the cross-population for the first time by integrating GSE144304 and GSE287726 two independent transcriptome datasets, clarified the central role of “metabolic-inflammatory-immune” synergistic disorder in the occurrence of frailty, and revealed highly consistent molecular characteristics in frailty. Based on this, a high-performance frailty diagnosis model is constructed. These findings provide new important insights for understanding the mechanisms of frailty research and developing clinical screening tools.

First, the most prominent finding of this study is the systemic dysregulation of vitamin metabolism pathways and ABC transporter pathways in frail individuals. Studies have shown^[10]^ that the abnormal function of ABC transporters as “molecular pumps” on cell membranes may affect the transmembrane transport of various nutrients, including vitamins^[11]^, thereby exacerbating cellular energy crises and dysfunction. In particular, vitamin D can also reverse the expression and function of some ABC transporters^[12]^. A study^[13]^ by Tachibana et al pointed out that the active form of vitamin D (1,25 (OH)_2_D_3_) can activate the transcription of the ABCB1 gene through the vitamin D receptor (VDR) and significantly increase the protein activity of ABCC2^[14]^. The function of vitamin D is far more than involved in the regulation of calcium and phosphorus homeostasis, and it plays a key role in maintaining muscle function, inhibiting inflammatory responses, and regulating the immune system^[15]^. Sarcopenia and immunocompromise, which are common in frail patients^[16]^, are likely to be directly related to vitamin D metabolism disorders^[17]^, a finding of far-reaching biological significance. In this study, it was found that ABCA1 and ABCG2 genes of the ABC transporter pathway were significantly down-regulated in frailty samples, and GSEA validation confirmed that this pathway was significantly enriched in both datasets. As a key transporter of vitamin D, the decreased expression of ABCG2 can lead to insufficient accumulation of active vitamin D (1,25-(OH) _2_ D _3_ ) in cells, thereby weakening the regulatory effect of vitamin D on immune inflammation^[10]^. At the same time, vitamin D deficiency may further down-regulate ABC transporter gene expression by inhibiting the VDR signaling pathway, forming a vicious cycle of “metabolism-transport”, which ultimately promotes the development of frailty.

Secondly, this study strongly confirms that chronic inflammation is a driver of frailty that cannot be ignored. Studies have shown that^[18]^ systemic chronic inflammatory states not only promote the progression of frailty, but also further increase the risk of adverse prognosis in frail populations. In both datasets, the co-activation of inflammation-related pathways such as cytokine-cytokine receptor interactions, complement-coagulation cascades, and NF-κB signaling pathway is highly consistent with the “inflammatory senescence” theory^[2]^.

The enrichment of cytokine signaling and cell adhesion molecules suggests that persistent and low-level activation of the innate immune system is an important driver of frailty^[2, 19]^. Overexpression of cytokines such as IL6 and TNF-α can lead to muscle loss and decreased neurological function^[4]^; Immune suppression (such as decreased T cell activity) weakens the body’s ability to eliminate inflammation and further aggravates tissue damage^[20]^. Other studies have added^[21]^ that there are also phenomena such as CD8+ and CCR5+ T cell expansion and weakened antibody response to influenza and other vaccines in frail populations^[22]^, and these immune inflammatory system disorders will lead to a decrease in the body’s resistance^[23]^, decline in physiological reserve function, and ultimately exacerbate frailty. Our innovation is to take these known assumptions to a more reliable level through repeated evidence from independent datasets.

At the level of translational medicine, the value of this study is significant. The SVM machine learning model^[24]^based on 612 intersecting DEGs achieved an AUC value of 0.959 in the independent validation set, demonstrating excellent diagnostic performance and generalization ability. This suggests that this set of signature genes excavated from the peripheral blood transcriptome provides candidate targets for the development of non-invasive or minimally invasive blood biomarker detection kits. This is expected to enable early identification and risk stratification of frailty in the future.

Of course, there are some limitations to this study. First, its cross-sectional study design cannot infer the causal direction of the discovered pathways, and whether these molecular changes are the cause, causality or concomitant phenomenon of frailty need to be verified by prospective cohort studies and animal models or cell experiments in the future. Second, despite the use of dual dataset validation, there is room for expansion in the total sample size (n=150), and the inclusion of more diverse and larger cohorts in the future will help further confirm the stability of these markers. Finally, transcriptome data are derived from peripheral blood, which is convenient for clinical translation, but may not fully reflect the molecular changes of key target tissues of frailty such as muscles and nerves.

## 5. Conclusions

In summary, we used bioinformatics integration analysis to reveal systemic disturbances in vitamin D metabolism, ABC transporter, and inflammatory signaling pathways in frail states. These molecular signatures validated across datasets not only deepen the understanding of frailty mechanisms, but also provide potential biomarkers and theoretical basis for early diagnosis and targeted intervention.

